# The Development of a Novel Nanobody Therapeutic for SARS-CoV-2

**DOI:** 10.1101/2020.11.17.386532

**Authors:** Gang Ye, Joseph P. Gallant, Christopher Massey, Ke Shi, Wanbo Tai, Jian Zheng, Abby E. Odle, Molly A. Vickers, Jian Shang, Yushun Wan, Aleksandra Drelich, Kempaiah R. Kempaiah, Vivian Tat, Stanley Perlman, Lanying Du, Chien-Te Tseng, Hideki Aihara, Aaron M. LeBeau, Fang Li

## Abstract

Combating the COVID-19 pandemic requires potent and low-cost therapeutics. We identified a novel series of single-domain antibodies (i.e., nanobody), Nanosota-1, from a camelid nanobody phage display library. Structural data showed that *Nanosota-1* bound to the oft-hidden receptor-binding domain (RBD) of SARS-CoV-2 spike protein, blocking out viral receptor ACE2. The lead drug possessing an Fc tag (*Nanosota-1C-Fc*) bound to SARS-CoV-2 RBD with a K_d_ of 15.7picomolar (∼3000 times more tightly than ACE2 did) and inhibited SARS-CoV-2 infection with an ND_50_ of 0.16microgram/milliliter (∼6000 times more potently than ACE2 did). Administered at a single dose, *Nanosota-1C-Fc* demonstrated preventive and therapeutic efficacy in hamsters subjected to SARS-CoV-2 infection. Unlike conventional antibody drugs, *Nanosota-1C-Fc* was produced at high yields in bacteria and had exceptional thermostability. Pharmacokinetic analysis of *Nanosota-1C-F*c documented a greater than 10-day *in vivo* half-life efficacy and high tissue bioavailability. *Nanosota-1C-Fc* is a potentially effective and realistic solution to the COVID-19 pandemic.

**Impact statement:** Potent and low-cost *Nanosota-1* drugs block SARS-CoV-2 infections both *in vitro* and *in vivo* and act both preventively and therapeutically.

## Introduction

The novel coronavirus SARS-CoV-2 has led to the COVID-19 pandemic, devastating human health and the global economy (*1, 2*). Anti-SARS-CoV-2 drugs are urgently needed to treat patients, save lives, and revive economies. Yet daunting challenges confront the development of such drugs. Though small molecule drugs could theoretically target SARS-CoV-2, they can take years to develop and their use is often limited by poor specificity and off-target effects. Repurposed drugs, developed against other viruses, also have low specificity against SARS-CoV-2. Therapeutic antibodies can be identified and generally have high specificity; however, their expression in mammalian cells often leads to low yields and high production costs (*3, 4*). A realistic therapeutic solution to COVID-19 must be potent and specific, yet easy to produce.

Nanobodies are unique antibodies derived from heavy chain-only antibodies found in members of the camelidae family (llamas, alpacas, camels, etc.) (Fig. S1) (*5, 6*). Because of their small size (2.5 nm by 4 nm; 12-15 kDa) and unique binding domains, nanobodies offer many advantages over conventional antibodies including the ability to bind cryptic epitopes on their antigen, high tissue permeability, ease of production and thermostability (*7, 8*). Although small, nanobodies bind their targets with high affinity and specificity due to an extended antigen-binding region (*7, 8*). Furthermore, it has been documented that they have low toxicity and immunogenicity in humans, if any (*7, 8*). One drawback of nanobodies is their quick clearance by kidneys due to their small size; this can be overcome by adding tags to increase the molecular weight to a desired level. Underscoring the potency and safety of nanobodies as human therapeutics, a nanobody drug was recently approved for clinical use in treating a blood clotting disorder (*9*). Additionally, due to their superior stability, nanobodies can be inhaled to treat lung diseases (*10*) or ingested to treat intestine diseases (*11*). Nanobodies are currently being developed against SARS-CoV-2 to combat COVID-19 (*12, 13*). However, to date, none of the reported nanobodies have been evaluated for therapeutic efficacy *in vivo*.

The receptor-binding domain (RBD) of the SARS-CoV-2 spike protein is a prime target for therapeutic development (*14*). The spike protein guides coronavirus entry into host cells by first binding to a receptor on the host cell surface and then fusing the viral and host membranes (*15, 16*). The RBDs of SARS-CoV-2 and a closely related SARS-CoV-1 both recognize human angiotensin-converting enzyme 2 (ACE2) as their receptor (*14, 17-19*). Previously, we showed that SARS-CoV-1 and SARS-CoV-2 RBDs both contain a core structure and a receptor-binding motif (RBM), and that SARS-CoV-2 RBD has significantly higher ACE2-binding affinity than SARS-CoV-1 RBD due to several structural changes in the RBM (*20, 21*). We further showed that SARS-CoV-2 RBD is more hidden than SARS-CoV-1 RBD in the entire spike protein as a possible viral strategy for immune evasion (*22*). Hence, to block SARS-CoV-2 binding to ACE2, a nanobody drug would need to bind to SARS-CoV-2 RBD more tightly than ACE2.

Here, we report the development of a novel series of anti-SARS-CoV-2 nanobody therapeutics, *Nanosota-1*. Identified by screening a camelid nanobody phage display library against the SARS-CoV-2 RBD, the *Nanosota-1* series bound potently to the SARS-CoV-2 RBD and were effective at inhibiting SARS-CoV-2 infection *in vitro*. The best performing drug, *Nanosota-1C-Fc*, demonstrated preventative and therapeutic efficacy in a hamster model of SARS-CoV-2 infection. *Nanosota-1C-Fc* was produced at high yields easily scalable for mass production and was also found to have a pharmacologically relevant *in vivo* half-life and excellent bioavailability. Our data suggest that *Nanosota-1c-Fc* may provide an effective solution to the COVID-19 pandemic.

## Results

### *Nanosota-1* was identified by phage display

For the rapid identification of virus-targeting nanobodies, we constructed a naïve nanobody phage display library using B cells isolated from the spleen, bone marrow, and blood of nearly a dozen non-immunized llamas and alpacas (Fig. 1). Recombinant SARS-CoV-2 RBD, expressed and purified from mammalian cells, was screened against the library to identify RBD-targeting nanobodies. Select nanobody clones were tested in a preliminary screen for their ability to neutralize SARS-CoV-2 pseudovirus entry into target cells (see below for more details about the assay). The nanobody that demonstrated the highest preliminary neutralization potency was named *Nanosota-1A* and then subjected to two rounds of affinity maturation. For each round of affinity maturation, random mutations were introduced to the whole gene of *Nanosota-1A* through error-prone PCR, and mutant phages were selected for enhanced binding to SARS-CoV-2 RBD. Nanobodies contain four framework regions (FRs) as structural scaffolds and three complementarity-determining regions (CDRs) for antigen binding. The nanobody after the first round of affinity maturation, named *Nanosota-1B*, possessed one mutation in CDR3 and two other mutations in FR3 (near CDR3). Affinity maturation of *Nanosota-1B* resulted in *Nanosota-1C*, which possessed one mutation in CDR2 and another mutation in FR2. We next made an Fc-tagged version of *Nanosota-1C*, termed *Nanosota-1C-Fc*, to create a bivalent construct with increased molecular weight.

**Figure 1:**
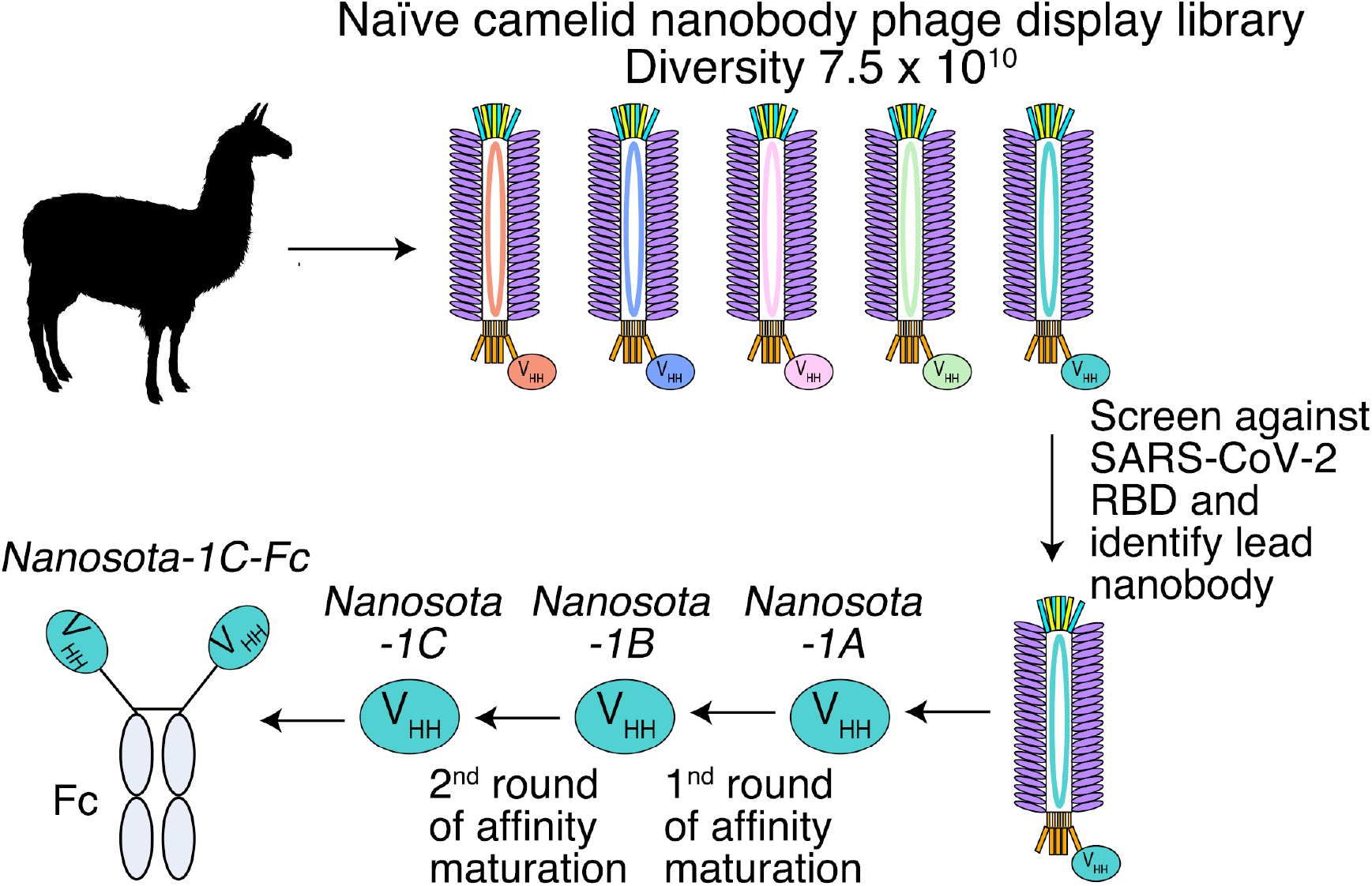
Construction of a camelid nanobody phage display library and use of this library for screening of anti-SARS-CoV-2 nanobodies. A large-sized (diversity 7.5 × 10^10^), naïve nanobody phage display library was constructed using B cells of over a dozen llamas and alpacas. Phages were screened for their high binding affinity for SARS-CoV-2 RBD. Nanobodies expressed from the selected phages were further screened for their potency in neutralizing SARS-CoV-2 pseudovirus entry. The best performing nanobody was subjected to two rounds of affinity maturation.

### *Nanosota-1* tightly bound to the SARS-CoV-2 RBD and completely blocked out ACE2

To understand the structural basis for the binding of *Nanosota-1* drugs to SARS-CoV-2 RBD, we determined the crystal structure of SARS-CoV-2 RBD complexed with *Nanosota-1C*. The structure showed that *Nanosota-1C* binds close to the center of the SARS-CoV-2 RBM (Fig. 2A). When the structures of the RBD/*Nanosota-1C* complex and the RBD/ACE2 complex were superimposed together, significant clashes occurred between ACE2 and *Nanosota-1C* (Fig. 2B), suggesting that *Nanosota-1C* binding to the RBD blocks ACE2 binding to the RBD. Moreover, trimeric SARS-CoV-2 spike protein is present in two different conformations: the RBD stands up in the open conformation but lies down in the closed conformation (*22-24*). When the structures of the RBD/*Nanosota-1C* complex and the closed spike were superimposed together, no clash was found between RBD-bound *Nanosota-1C* and the rest of the spike protein (Fig. S2A). In contrast, severe clashes were identified between RBD-bound ACE2 and the rest of the spike protein in the closed conformation (Fig. S2B). Additionally, neither RBD-bound *Nanosota-1C* nor RBD-bound ACE2 had clashes with the rest of the spike protein in the open conformation (Fig. S2C, S2D). Thus, *Nanosota-1C* can access the spike protein in both its open and closed conformations, whereas ACE2 can only access the spike protein in its closed conformation. Overall, our structural data reveal that *Nanosota-1C* is an ideal RBD-targeting drug that not only blocks virus binding to its receptor, but also accesses its target in the spike protein in different conformations.

**Figure 2:**
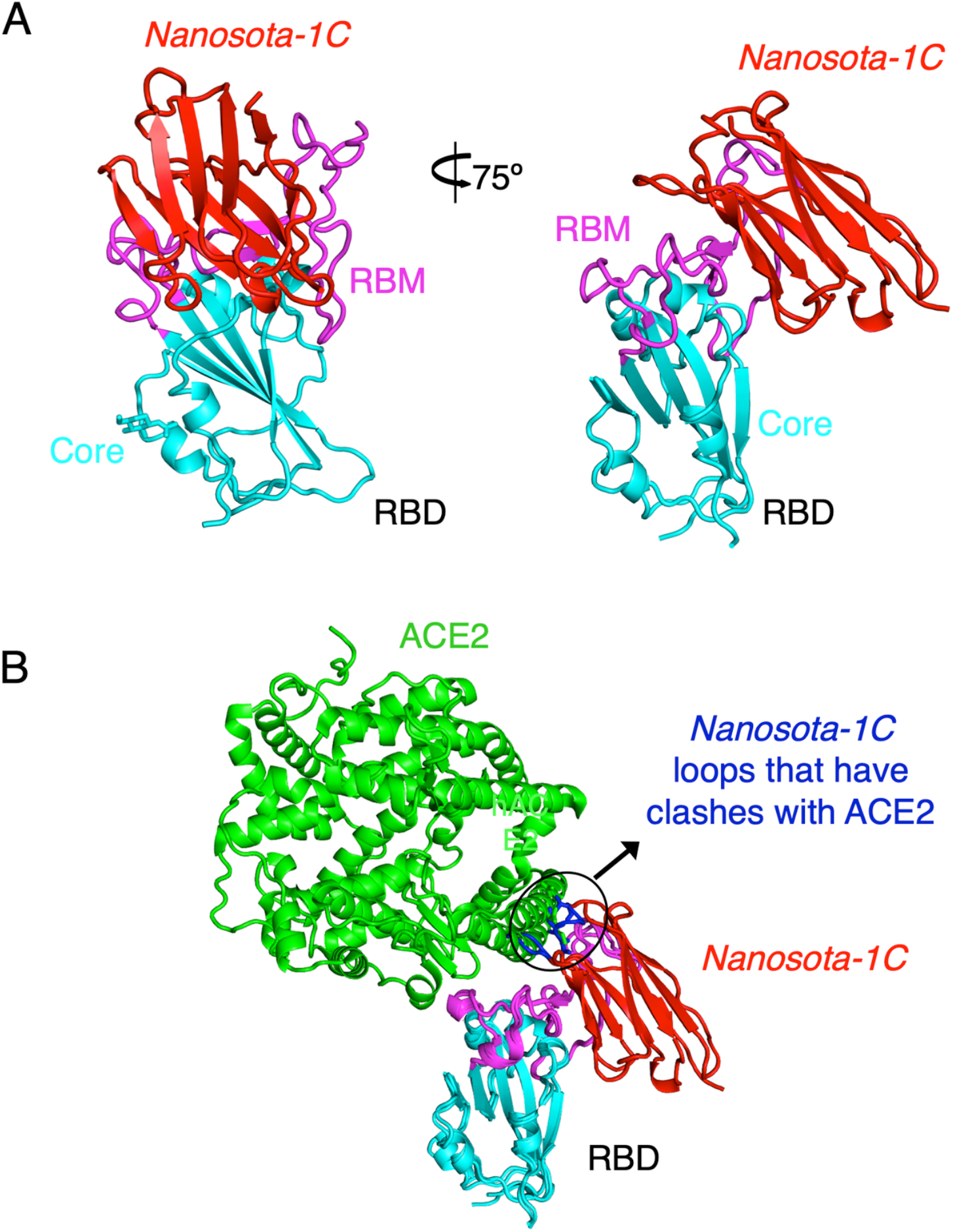
Crystal structure of SARS-CoV-2 RBD complexed with *Nanosota-1C*. (A) Structure of SARS-CoV-2 RBD complexed with *Nanosota-1C*, viewed at two different angles. *Nanosota-1C* is in red, the core structure of RBD is in cyan, and the receptor-binding motif (RBM) of RBD is in magenta. (B) Overlay of the structures of the RBD/*Nanosota-1C* complex and RBD/ACE2 complex (PDB 6M0J). ACE2 is in green. The structures of the two complexes were superimposed based on their common RBD structure. The *Nanosota-1C* loops that have clashes with ACE2 are in blue.

To corroborate our structural data on the *Nanosota-1*/ACE2 interactions, we performed binding experiments between *Nanosota-1* drugs and SARS-CoV-2 RBD using recombinant ACE2 for comparison. The binding affinity between the nanobodies and the RBD were measured by surface plasmon resonance (Table 1; Fig. S3). *Nanosota-1A, −1B, and −1C* bound to the RBD with increasing affinity (K_d_ - from 228 nM to 14 nM), confirming success of the stepwise affinity maturation. *Nanosota-1C-Fc* had the highest RBD-binding affinity (K_d_ −15.7 pM), which was ∼3,000 times tighter than the RBD-binding affinity of ACE2. Moreover, compared with ACE2, *Nanosota-1C-Fc* bound to the RBD with a higher *k*_*on*_ and a lower *k*_*off*_, demonstrating significantly faster binding and slower dissociation. Next, we investigated the competitive binding among *Nanosota-1C*, ACE2, and RBD using protein pull-down assay (Fig. S4A). ACE2 and *Nanosota-1C* were mixed together in different ratios in solution, with the concentration of ACE2 kept constant; RBD-Fc was added to pull down ACE2 and *Nanosota-1C* from solution. The result showed that as the concentration of *Nanosota-1C* increased, less ACE2 was pulled down by the RBD. Thus, ACE2 and *Nanosota-1C* bound competitively to the RBD. We then analyzed the competitive binding using gel filtration chromatography (Fig. S4B). ACE2, *Nanosota-1C*, and RBD were mixed, with both ACE2 and *Nanosota-1C* in molar excess over the RBD. Analysis by gel filtration chromatography documented that no ternary complex of ACE2, *Nanosota-1C*, and RBD formed; instead, only binary complexes of RBD/ACE2 and RBD/*Nanosota-1C* were detected. Hence, the bindings of ACE2 and *Nanosota-1C* to the RBD are mutually exclusive.

**Table 1.**
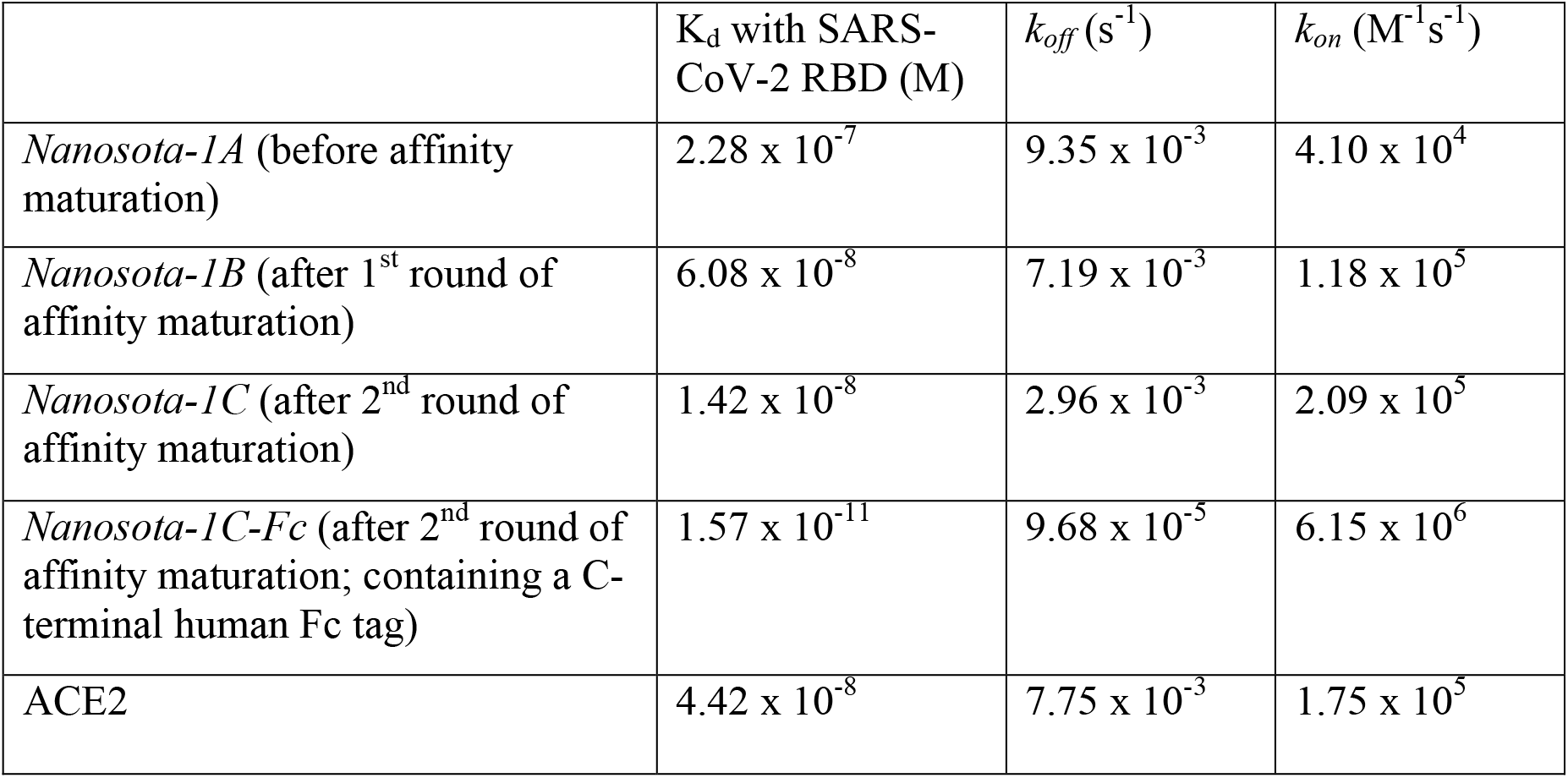
Binding affinities between *Nanosota-1* drugs and SARS-CoV-2 RBD as measured using surface plasmon resonance. The previously determined binding affinity between human ACE2 and RBD is shown as a comparison (*20*).

### *Nanosota-1C-Fc* potently neutralized SARS-CoV-2 infection *in vitro* and *in vivo*

The ability of the *Nanosota-1* drugs to neutralize SARS-CoV-2 infection *in vitro* was investigated nex. Both a SARS-CoV-2 pseudovirus entry assay and authentic SARS-CoV-2 infection assay were performed (Fig. 3). For the pseudovirus entry assay, retroviruses pseudotyped with SARS-CoV-2 spike protein (i.e., SARS-CoV-2 pseudoviruses) were used to enter human ACE2-expressing HEK293T cells in the presence of an inhibitor. The efficacy of the inhibitor was expressed as the concentration capable of neutralizing 50% of the entry efficiency (i.e., 50% Neutralizing Dose or ND_50_). *Nanosota-1C-Fc* had an ND_50_ for the SARS-CoV-2 pseudovirus of 0.27 µg/ml, which was ∼10 times more potent than monovalent *Nanosota-1C* (2.52 µg/ml) and over 100 times more potent than ACE2 (44.8 µg/ml) (Fig. 3A). Additionally, *Nanosota-1* drugs potently neutralized SARS-CoV-2 pseudovirus bearing the D614G mutation in the SARS-CoV-2 spike protein (Fig. S5), which has become prevalent in many strains (*25*). For the authentic virus infection assay, live SARS-CoV-2 was used to infect Vero cells in the presence of an inhibitor. Efficacy of the inhibitor was described as the concentration capable of reducing the number of virus plaques by 50% (i.e., ND_50_). *Nanosota-1C-Fc* had an ND_50_ of 0.16 µg/ml, which was ∼20 times more potent than monovalent *Nanosota-1C* (3.23 µg/ml) and ∼6000 times more potent than ACE2 (980 µg/ml) (Fig. 3B; Fig. S6). Overall, both *Nanosota-1C-Fc* and *Nanosota-1C* are potent inhibitors of SARS-CoV-2 pseudovirus entry and authentic SARS-CoV-2 infection.

**Figure 3.**
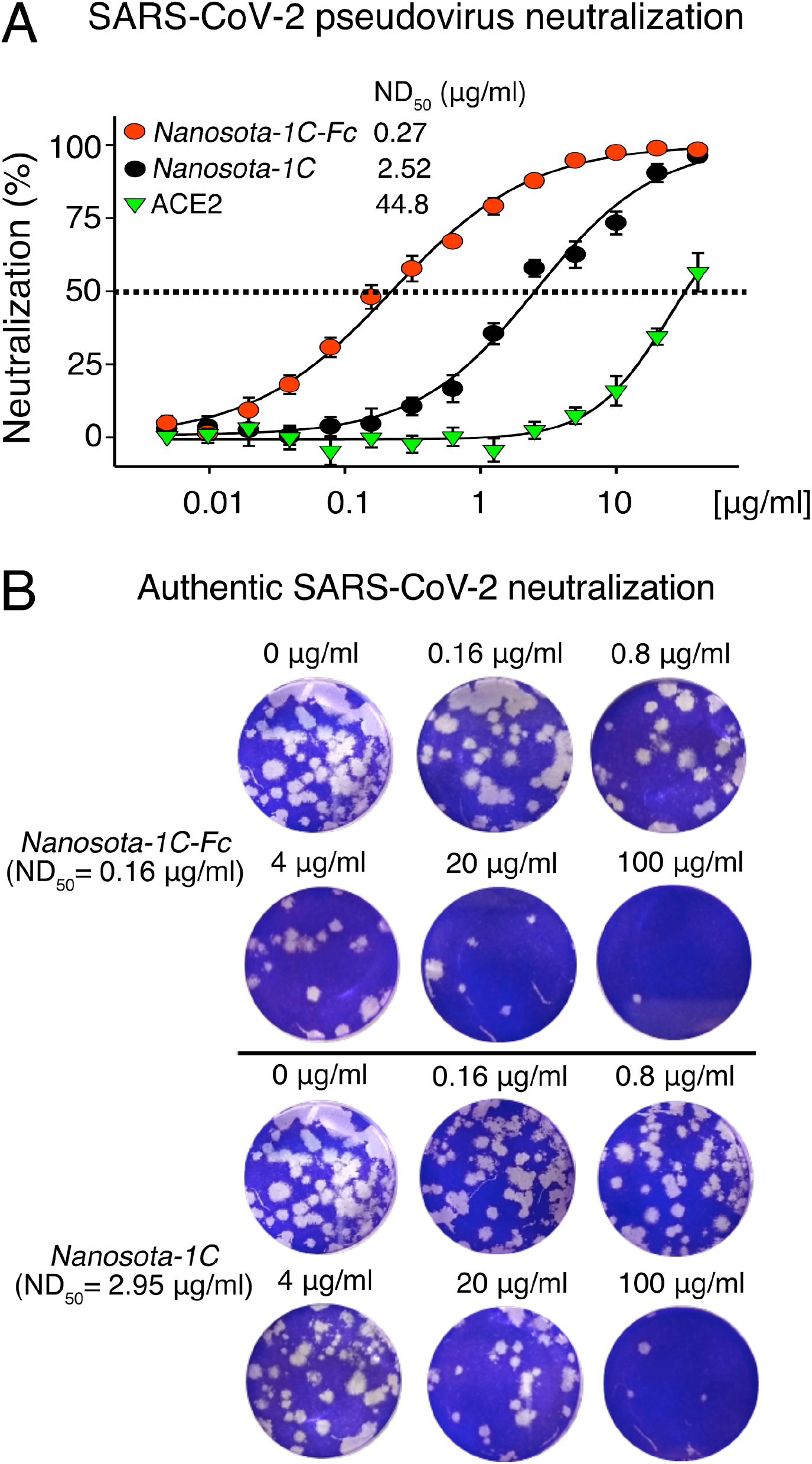
Efficacy of *Nanosota-1* drugs in neutralizing SARS-CoV-2 infections *in vitro*. (A) Neutralization of SARS-CoV-2 pseudovirus entry into target cells by one of three inhibitors: *Nanosota-1C-Fc, Nanosota-1C*, and recombinant human ACE2. Retroviruses pseudotyped with SARS-CoV-2 spike protein (i.e., SARS-CoV-2 pseudoviruses) were used to enter HEK293T cells expressing human ACE2 in the presence of the inhibitor at various concentrations. Entry efficiency was characterized via a luciferase signal indicating successful cell entry. Data are the mean ± SEM (n = 4). Nonlinear regression was performed using a log (inhibitor) versus normalized response curve and a variable slope model (R^2^ > 0.95 for all curves). The efficacy of each inhibitor was expressed as the 50% Neutralizing Dose or ND_50_. The assay was repeated three times (biological replication: new aliquots of pseudoviruses and cells were used for each repeat). (B) Neutralization of authentic SARS-CoV-2 infection of target cells by one of two inhibitors: *Nanosota-1C-Fc* and *Nanosota-1C*. The potency of *Nanosota-1* drugs in neutralizing authentic SARS-CoV-2 infections was evaluated using a SARS-CoV-2 plaque reduction neutralization test (PRNT) assay. 80 pfu infectious SARS-CoV-2 particles were used to infect Vero E6 cells in the presence of the inhibitor at various concentrations. Infection was characterized as the number of virus plaques formed in overlaid cells. Images of virus plaques for each inhibitor at the indicated concentrations are shown. Each image represents data from triplications. The efficacy of each inhibitor was calculated and expressed as the concentration capable of reducing the number of virus plaques by 50% (i.e., ND_50_). The assay was repeated twice (biological replication: new aliquots of virus particles and cells were used for each repeat).

After the *in vitro* studies, we next evaluated the therapeutic efficacy of the lead drug *Nanosota-1C-Fc* in a hamster model challenged with SARS-CoV-2 via intranasal inoculation. In addition to an untreated control group, three groups of animals were injected with a single dose of *Nanosota-1C-Fc*: (i) 24 hours pre-challenge at 20 mg/kg body weight, (ii) 4 hours post-challenge at 20 mg/kg, and (iii) 4 hours post-challenge at 10 mg/kg. As previously validated in this model (*26*), body weight, tissue pathology and virus titers in nasal swabs were used as metrics of therapeutic efficacy. In the untreated control group, weight loss was precipitously starting on day 1 post-challenge with the lowest weight recorded on day 6 (Fig. 4A). Nasal virus titers were high on day 1 and remained high on day 5 before a decline (Fig. S7). Pathology analysis on tissues collected on day 10 revealed moderate hyperplasia in the bronchial tubes (i.e., bronchioloalveolar hyperplasia) (Fig. 4B), with little hyperplasia in the lungs. These data are consistent with previous reports showing that SARS-CoV-2 mainly infects the nasal mucosa and bronchial epithelial cells of this hamster model (*26*). In contrast, hamsters that received *Nanosota-1C-Fc* 24-hours pre-challenge were protected from SARS-CoV-2, as evidenced by the metrics of no weight loss, no bronchioloalveolar hyperplasia, and significantly reduced nasal virus titers (Fig. 4, Fig. S7). When administered 4 hours post-challenge, *Nanosota-1C-Fc* also effectively protected hamsters from SARS-CoV-2 infections at either dosage (20 or 10 mg/kg), as evidenced by the favorable therapeutic metrics (Fig. 4, Fig. S7). Overall, *Nanosota-1C-Fc* was effective at combating SARS-CoV-2 infections both preventively and therapeutically.

**Figure 4.**
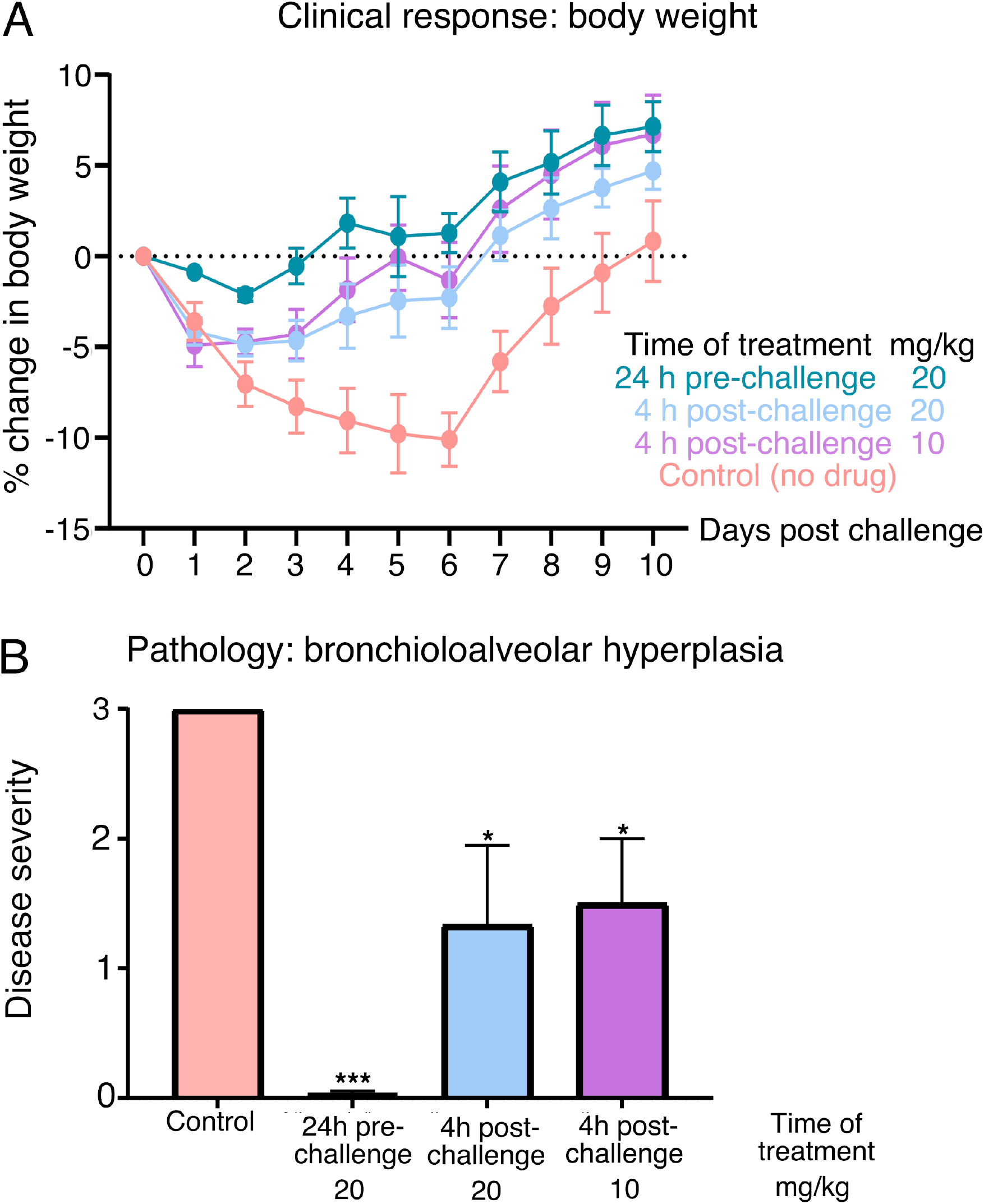
Efficacy of *Nanosota-1* drugs in protecting hamsters from SARS-CoV-2 infections. Hamsters (6 per group) were injected with a single dose of *Nanosota-1C-Fc* at the indicated time point and the indicated dosage. At day 0 all groups (experimental and control) were challenged with SARS-CoV-2 (at a titer of 10^6^ Median Tissue Culture Infectious Dose or TCID_50_). (A) Body weights of hamsters were monitored on each day and percent change in body weight relative to day 0 was calculated for each hamster. Data are the mean ± SEM (n = 6). ANOVA on group as a between-group factor and day (1-10) as a within-group factor revealed significant differences between the control group and each of the following groups: 24 hour pre-challenge (20 mg/kg) group (F(1, 10) = 17.80, *p* = .002; effect size *η*_*p*_^2^ = .64), 4 hour post-challenge (20 mg/kg) group (*F*(1, 10) = 5.02, *p* = .035; *η*_*p*_^2^ = .37), and 4 hour post-challenge (10 mg/kg) group (*F*(1, 10) = 7.04, *p* = .024, *η*_*p*_^2^ = .41). All *p-*values are two-tailed. (B) Tissues of bronchial tubes from each of the hamsters were collected on day 10 and scored for the severity of bronchioloalveolar hyperplasia: 3 - moderate; 2 - mild; 1 - minimum; 0 - none. Data are the mean ± SEM (n = 6). A comparison between the control group and each of other groups was performed using one-tailed Student’s t-test for directional tests. \*\*\**p* < 0.001; \**p* < 0.05.

### *Nanosota-1C-Fc* is stable *in vitro* and *in vivo* with excellent bioavailability

With the lead drug *Nanosota-1C-Fc* demonstrating therapeutic efficacy *in vivo*, we characterized other parameters important to its clinical translation. First, we expressed *Nanosota-1C-Fc* in bacteria for all the experiments carried out in the current study (Fig. 5A). After purification on protein A column and gel filtration, the purity of *Nanosota-1C-Fc* was nearly 100%. With no optimization, the expression yield reached 40 mg/L of bacterial culture. Second, we investigated the *in vitro* stability of *Nanosota-1C-Fc* incubated at four temperatures (−80°C, 4°C, 25°C or 37°C) for one week and then measured the remaining SARS-CoV-2 RBD-binding capacity by ELISA (Fig. 5B). With −80°C as a baseline, *Nanosota-1C-Fc* retained nearly all of its RBD-binding capacity at the temperatures surveyed. Third, we measured the *in vivo* stability of *Nanosota-1C-Fc* (Fig. 5C). *Nanosota-1C-Fc* was injected into mice via tail vein. Sera were obtained at different time points and measured for their SARS-CoV-2 RBD-binding capacity by ELISA. *Nanosota-1C-Fc* retained most of its RBD-binding capability after 10 days *in vivo*. Antithetically, *Nanosota-1C* was stable for only several hours *in vivo*. (Fig. S8A). Last, we examined the biodistribution of *Nanosota-1C-Fc* in mice (Fig. 5D). *Nanosota-1C-Fc* was radiolabeled with zirocinium-89 and injected systemically into mice. Tissues were collected at various time points and biodistribution of *Nanosota-1C-Fc* was quantified by scintillation counter. After three days, *Nanosota-1C-Fc* remained at high levels in the blood, lung, heart, kidney, liver and spleen, all of which are targets for SARS-CoV-2 (*27*); moreover, it remained at low levels in the intestine, muscle and bones. In contrast, *Nanosota-1C* had poor biodistribution documenting high renal clearance (Fig. S8B). Overall, our findings suggest that *Nanosota-1C-Fc* is potent SARS-CoV-2 therapeutic with translational values applicable to the world’s vast population.

**Figure 5.**
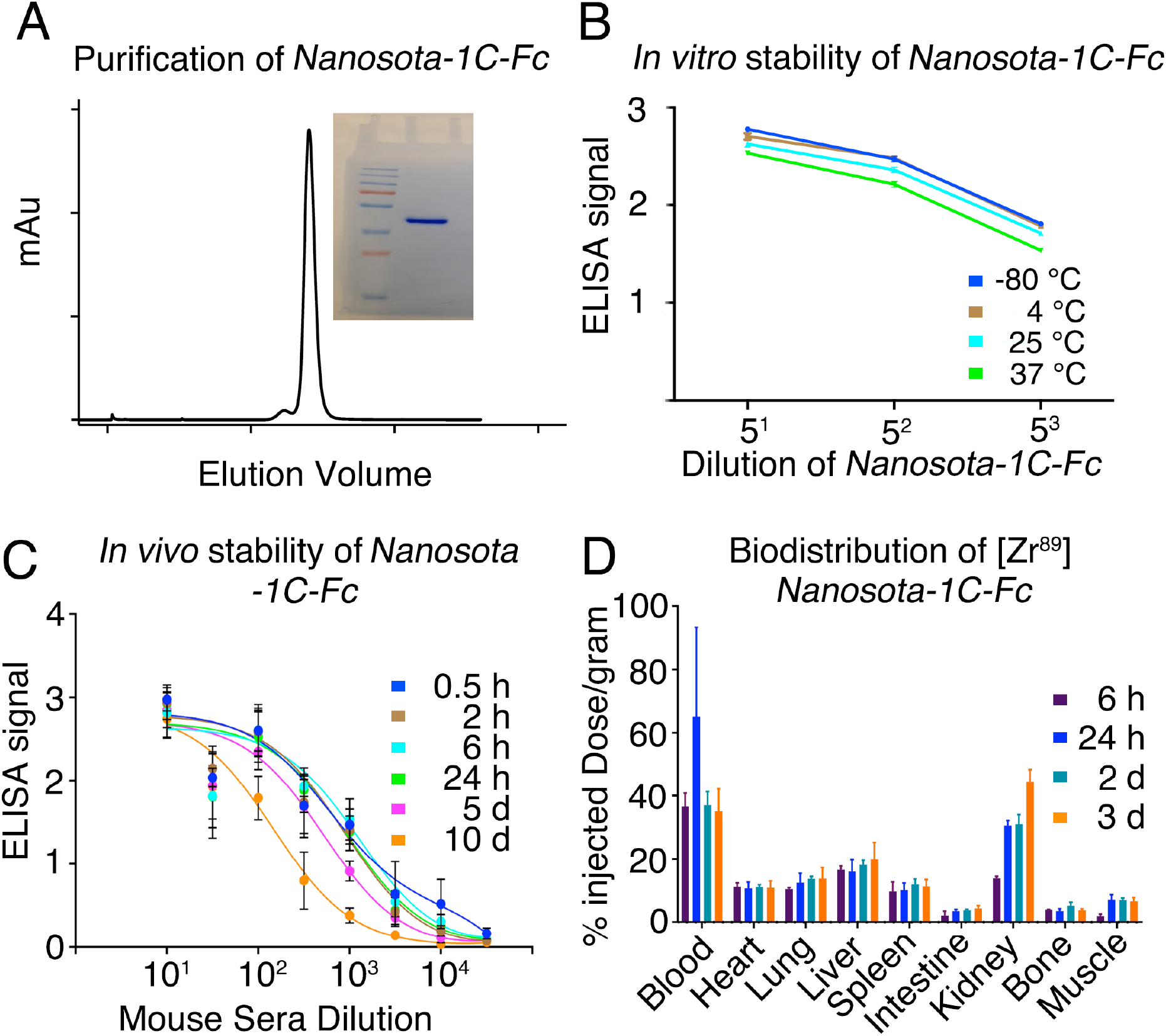
Analysis of expression, purification and pharmacokinetics of *Nanosota-1C-Fc*. (A) Purification of *Nanosota-1C-Fc* from bacteria. The protein was nearly 100% pure after gel filtration chromatography, as demonstrated by its elution profile and SDS-PAGE (stained by Coomassie blue). The yield of the protein was 40 mg/L of bacterial culture, without any optimization of the expression. (B) *In vitro* stability of *Nanosota-1C-Fc*. The protein was stored at indicated temperatures for a week, and then a dilution ELISA was performed to evaluate its SARS-CoV-2 RBD-binding capability. Data are the mean ± SEM (n = 4). (C) *In vivo* stability of *Nanosota-1C-Fc. Nanosota-1C-Fc* was injected into mice, mouse sera were collected at different time points, and *Nanosota-1C-Fc* remaining in the sera was detected for its SARS-CoV-2 RBD-binding capability as displayed in a dilution ELISA. Data are the mean ± SEM (n = 3). (D) Biodistribution of [^89^Zr]Zr-*Nanosota-1C-Fc. Nanosota-1C-Fc* was radioactively labeled with ^89^Zr and injected into mice via tail vein injection. Different tissues or organs were collected at various time points (n=3 mice per time point). The amount of *Nanosota-1C-Fc* present in each tissue or organ was measured through examining the radioactive count of each tissue or organ. Data are the mean ± SEM (n = 3).

## Discussion

Nanobody therapeutics derived from camelid antibodies potentially offer a realistic solution to the COVID-19 pandemic compared to conventional antibodies. Currently, there have only been a few reports of nanobody drugs that specifically target SARS-CoV-2 (*12, 13*). Those reported were developed against SARS-CoV-2 RBD, either blocking out ACE2 or locking the RBD in the closed inactive state on the spike protein (*12, 13*). None of the nanobodies have been evaluated in animal models for their anti-SARS-CoV-2 therapeutic efficacy. From our novel library, we developed a series of nanobody drugs, named *Nanosota-1*, that specifically target the SARS-CoV-2 RBD. Two rounds of affinity maturation yielded *Nanosota-1C* which bound to the RBD with high affinity. Addition of an Fc tag to make a bivalent construct with increased molecular weight and picomolar RBD-binding affinity resulted in the best performing drug *Nanosota-1C-Fc*. Our structural and biochemical data showed that binding of *Nanosota-1C* to the RBD blocked virus binding to viral receptor ACE2. A unique feature of the SARS-CoV-2 spike protein is that it is present in two different conformations, an RBD-up open conformation for receptor binding and an RBD-down closed conformation for immune evasion (20, 22, 23). Due to its small size as well as its ideal binding site on the RBD, *Nanosota-1* can bind to the spike protein in both conformations. In contrast, ACE2 can only bind to the spike protein in its open conformation. Thus, *Nanosota-1* drugs are ideal RBD-targeting therapeutics - they can chase down and inhibit SARS-CoV-2 viral particles whether they are infecting cells or hiding from immune surveillance. As a result of this unique property, both *Nanosota-1C* and *Nanosota-1C-Fc* exhibited a profound therapeutic effect *in vitro* against SARS-CoV-2 pseudovirus and authentic SARS-CoV-2. *Nanosota-1C-Fc* was also found to be the first anti-SARS-CoV-2 camelid nanobody-based therapeutic reported in the literature to demonstrate efficacy in an animal model. Additionally, *Nanosota-1C-Fc* was the first anti-SARS-CoV-2 nanobody to have been characterized for ease of production and purification, *in vitro* and *in vivo* stabilities, and biodistribution. These features are critical for the implementation of *Nanosota-1C-F*c as a COVID-19 therapeutic.

When evaluating the anti-SARS-CoV-2 potency of the nanobody therapeutics, we used recombinant ACE2 as a comparison. Recombinant ACE2 was selected because *Nanosota-1* series directly compete with cell-surface ACE2 for the same binding site on the RBD. Our study showed that compared with ACE2, the best performing drug *Nanosota-1C-Fc* bound to the RBD ∼3000 fold more strongly, blocking out ACE2 binding to the RBD. Furthermore, compared with ACE2, *Nanosota-1C-Fc* inhibited SARS-CoV-2 pseudovirus entry ∼100 fold more effectively and inhibited authentic SARS-CoV-2 infections ∼6000 fold more effectively. Note that recombinant ACE2 has been shown to be a potent anti-SARS-CoV-2 inhibitor (*28*) and is currently undergoing clinical trials in Europe as an anti-COVID-19 drug. Compared with ACE2, the much higher anti-SARS-CoV-2 potency of *Nanosota-1C-Fc* was due to both its much higher RBD-binding affinity and its better access to the oft-hidden RBD in the spike protein. As a result, *Nanosota-1C-Fc* was a potent therapeutic *in vivo*. Remarkably, a single dose of *Nanosota-1C-Fc* effectively prevented SARS-CoV-2 infection in hamsters and also effectively treated SARS-CoV-2 infection in the same model. The hamster model is one of the best non-primate models available for studying anti-SARS-CoV-2 therapeutic efficacy, but it is limited by a short virus infection window; hence, repeated dosing was not evaluated. As a result, we were only able to dose the mice once via intraperitoneal injection. Because SARS-CoV-2 is fast acting in hamsters, the time points and dosages for drug administration in hamsters are difficult to directly translate to humans. Our supporting data document that *Nanosota-1c-Fc* is easy to produce in bacteria and has excellent bioavailability and pharmacokinetics when administered intravenously in mice. This suggests that *Nanosota-1C-Fc* may have therapeutic potential when administered intraperitoneal, intravenous or even intramuscular. These parameters will need to be determined in future studies in anticipation of clinical trials. Overall, *Nanosota-1C-Fc* has proven to be an effective therapeutic in the model that we currently have available.

How can the novel nanobody therapeutics help to end the COVID-19 pandemic? First, as evidence by our animal study, *Nanosota-1C-Fc* can be used to prevent SARS-CoV-2 infection. Because of its long *in vivo* half-life (>10 days), a single injected dose of *Nanosota-1C-Fc* can theoretically protect a person from SARS-CoV-2 infection for days or weeks in the outpatient setting, reducing the spread of SARS-CoV-2 in human populations. Second, we also learned from our *in vivo* study that *Nanosota-1C-Fc* can potentially be used to treat SARS-CoV-2 infections, thus, saving lives and alleviating symptoms in infected patients in the clinical setting. Third, though ephemeral in nature given its short half-life and rapid clearance from the blood, *Nanosota-1C* could be used as an inhaler to treat infections in the respiratory tracts (*10*) or as an oral drug to treat infections in the intestines (*11*). Overall, the novel series of *Nanosota-1* therapeutics can help minimize the mortality and morbidity of SARS-CoV-2 infections and help restore the economy and daily human activities. Given the wide distribution of SARS-CoV-2 in the world, large quantities of anti-SARS-CoV-2 therapeutics would need to be manufactured to provide for the world’s populations. This is only feasible with easy to produce and scalable molecules, such as *Nanosota-1* drugs, that are produced at high yields and have long *in vitro* and *in vivo* half-life. Therefore, if further validated in clinical trials, *Nanosota-1* therapeutics can provide a realistic and effective solution to help end the COVID-19 global pandemic.

## Acknowledgements

The development of *Nanosota-1* drugs and the animal testing were supported by funding from the University of Minnesota (to F.L.), NIH grants R01AI157975 (to F.L., A.M.L., L.D., S.P.), R01AI089728 (to F.L.), and R35GM118047 (to H.A.). A.M.L. is a 2013 Prostate Cancer Foundation Young Investigator and recipient of a 2018 Prostate Cancer Foundation Challenge Award. Experimental Pathology Laboratories analyzed pathology data on SARS-CoV-2-challenged hamsters. Crystallization screening was performed at Hauptman-Woodward Medical Research Institute and supported by NSF grant 2029943. X-ray diffraction data were collected at Advanced Photon Source beamline 24-ID-E and we thank Surajit Bannerjee for help in X-ray data collection. The University of Minnesota has filed a patent on Nanosota-1 drugs with F.L, G.Y., A.M.L., J.P.G., J.S., and Y.W. as inventors. We thank Professor Yuhong Jiang for consultation on the design of animal testing and statistical analysis and for editing the manuscript. Coordinates and structure factors have been deposited to the Protein Data Bank with accession number XXXX.

## Methods

### Ethics statement

This study was performed in strict accordance with the recommendations in the Guide for the Care and Use of Laboratory Animals of the National Institutes of Health. All of the animals were handled according to approved institutional animal care and use committee (IACUC) protocols of the University of Texas Medical Branch (protocol number 2007072) and of the University of Minnesota (protocol number 2009-38426A).

### Cell lines, plasmids and virus

HEK293T cells (American Type Culture Collection) were cultured in Dulbecco’s modified Eagle medium (DMEM) supplemented with 10% fetal bovine serum, 2 mM L-glutamine, 100 units/mL penicillin, and 100 µg/mL streptomycin (Life Technologies). ss320 *E. coli* (Lucigen), TG1 *E. coli* (Lucigen), SHuffle T7 *E. coli* (New England Biolabs) were grown in TB medium or 2YT medium with 100 mg/L ampicillin. Vero E6 cells (American Type Culture Collection) were grown in Eagle’s minimal essential medium (EMEM) supplemented with penicillin (100 units/ml), streptomycin (100 µg/ml), and 10% fetal bovine serum (FBS). SARS-CoV-2 spike (GenBank accession number QHD43416.1) and ACE2 (GenBank accession number NM_021804) were described previously (*20*). SARS-CoV-2 RBD (residues 319-529) was subcloned into Lenti-CMV vector (Vigene Biosciences) with an N-terminal tissue plasminogen activator (tPA) signal peptide and a C-terminal human IgG4 Fc tag or His tag. The ACE2 ectodomain (residues 1–615) was constructed in the same way except that its own signal peptide was used. *Nanosota-1A*, -*1B* and -*1C* were each cloned into PADL22c vector (Lucigen) with a N-terminal PelB leader sequence and C-terminal His tag and HA tag. *Nanosota-1C-Fc* was cloned into pET42b vector (Novagen) with a C-terminal human IgG_1_ Fc tag. SARS-CoV-2 (US_WA-1 isolate) from CDC (Atlanta) was used throughout the study. All experiments involving infectious SARS-CoV-2 were conducted at the University of Texas Medical Branch and University of Iowa in approved biosafety level 3 laboratories.

### Construction of camelid nanobody phage display library

The camelid nanobody phage display library was constructed as previously described (*29, 30*). Briefly, total mRNA was isolated from B cells from the spleen, bone marrow and blood of over a dozen non-immunized llamas and alpacas. cDNA was prepared from the mRNA. The cDNA was then used in nested PCR reactions to construct the DNA for the library. The first PCR reaction was to amplify the gene fragments encoding the variable domain of the nanobody. The second PCR reaction (PCR2) was used to add restriction sites (SFI-I), a PelB leader sequence, a His_6_ tag, and a HA tag. The PCR2 product was digested with SFI-I (New England Biolabs) and then was ligated with SFI-I-digested PADL22c vector. The ligated product was transformed via electroporation into TG1 *E. coli* (Lucigen). Aliquots of cells were spread onto 2YT agar plates supplemented with ampicillin and glucose, incubated at 30°C overnight, and then scraped into 2YT media. After centrifugation, the cell pellet was suspended into 50% glycerol and stored at −80°C. The library size was 7. 5 × 10^10^. To display nanobodies on phages, aliquots of the TG1 *E. coli* bank were inoculated into 2YT media, grown to early logarithmic phase, and infected with M13K07 helper phage.

### Camelid nanobody library screening

The above camelid nanobody phage display library was used in the bio-panning as previously described (*31*). Briefly, four rounds of panning were performed to obtain the SARS-CoV-2 RBD-targeting nanobodies with high RBD-binding affinity. The amounts of the RBD antigen used in coating the immune tubes in each round were 75 µg, 50 µg, 25 µg, and 10 µg, respectively. The retained phages were eluted using 1 ml 100 mM triethylamine and neutralized with 500 µl 1 M Tris-HCl pH 7.5. The eluted phages were amplified in TG1 *E. coli* and rescued with M13K07 helper phage. The eluted phages from round 4 were used to infect ss320 *E. coli*. Single colonies were picked into 2YT media and nanobody expressions were induced with 1 mM IPTG. The supernatants were subjected to ELISA for selection of strong binders (described below). The strong binders were then expressed and purified (described below) and subjected to SARS-CoV-2 pseudovirus entry assay for selection of anti-SARS-CoV-2 efficacy (described below). The lead nanobody after initial screening was named *Nanosota-1A*.

### Affinity maturation

Affinity maturation of *Nanosota-1A* was performed as previously described (*32*). Briefly, mutations were introduced into the whole gene of *Nanosota-1A* using error-prone PCR. Two rounds of error-prone PCR were performed using the GeneMorph II Random Mutagenesis Kit (Agilent Technologies). The PCR product was cloned into the PADL22c vector and transformed via electroporation into the TG1 *E. coli*. The library size was 6 × 10_8_. Three rounds of bio-panning were performed using 25 ng, 10 ng and 2 ng RBD-Fc, respectively. The strongest binder after affinity maturation was named *Nanosota-1B*. A second round of affinity maturation was performed in the same way as the first round, except that three rounds of bio-panning were performed using 10 ng, 2 ng and 0.5 ng RBD-Fc, respectively. The strongest binder after the second round of affinity maturation was named *Nanosota-1C*.

### Production of Nanosota-1 drugs

*Nanosota-1A, 1B* and *1C* were each purified from the periplasm of ss320 *E. coli* after the cells were induced by 1 mM IPTG. The cells were collected and re-suspended in 15 ml TES buffer (0.2 M Tris pH 8, 0.5 mM EDTA, 0.5 M sucrose), shaken on ice for 1 hour and then incubated with 40 ml TES buffer followed by shaking on ice for another hour. The protein in the supernatant was sequentially purified using a Ni-NTA column and a Superdex200 gel filtration column (GE Healthcare) as previously described (*20*). *Nanosota-1C-Fc* was purified from the cytoplasm of Shuffle T7 *E. coli*. The induction of protein expression was the same as above. After induction, the cells were collected, re-suspected in PBS and disrupted using Branson Digital Sonifier (Thermofisher). The protein in the supernatant was sequentially purified on protein A column and Superdex200 gel filtration column as previously described (*20*).

### Production of SARS-CoV-2 RBD and ACE2

HEK293T cells stably expressing SARS-CoV-2 RBD (containing a C-terminal His tag or Fc tag) or human ACE2 ectodomain (containing a C-terminal His tag) were made according to the E and F sections of the pLKO.1 Protocol from Addgene (http://www.addgene.org/protocols/plko/). The proteins were secreted to cell culture media, harvested, and purified on either Ni-NTA column (for His-tagged proteins) or protein A column (for Fc-tagged protein) and then on Superdex200 gel filtration column as previously described (*20*).

### ELISA

ELISA was performed to detect the binding between SARS-CoV-2 RBD and *Nanosota-1* drugs (either purified recombinant drugs or drugs in the mouse serum) as previously described (*33*). Briefly, ELISA plates were coated with recombinant SARS-CoV-2 RBD-His or RBD-Fc, and were then incubated sequentially with nanobody drugs, HRP-conjugated anti-llama antibody (1:5,000) (Sigma) or HRP-conjugated anti-human-Fc antibody (1:5,000) (Jackson ImmunoResearch). ELISA substrate (Invitrogen) was added to the plates, and the reactions were stopped with 1N H_2_SO4. The absorbance at 450 nm (A_450_) was measured using a Synergy LX Multi-Mode Reader (BioTek).

### Determination of the structure of SARS-CoV-2 RBD complexed with Nanosota-1C

To prepare the RBD/*Nanosota-1C* complex for crystallization, the two proteins were mixed together in solution and purified using a Superdex200 gel filtration column (GE Healthcare). The complex was concentrated to 10 mg/ml in buffer 20 mM Tris pH 7.2 and 200 mM NaCl. Crystals were screened at High-Throughput Crystallization Screening Center (Hauptman-Woodward Medical Research Institute) as previously described (*34*), and were grown in sitting drops at room temperature over wells containing 50 mM MnCl_2_, 50 mM MES pH 6.0, 20% (W/V) PEG 4000. Crystals were soaked briefly in 50 mM MnCl_2_, 50 mM MES pH 6.0, 25% (W/V) PEG 4000 and 30% ethylene glycol before being flash-frozen in liquid nitrogen. X-ray diffraction data were collected at the Advanced Photon Source beamline 24-ID-E. The structure was determined by molecular replacement using the structures of SARS-CoV-2 RBD (PDB 6M0J) and another nanobody (PDB 6QX4) as the search templates. Structure data and refinement statistics are shown in Table S1.

### Surface plasmon resonance assay

Surface plasmon resonance assay using a Biacore S200 system (GE Healthcare) was carried out as previously described (*20*). Briefly, SARS2-CoV-2 RBD-His was immobilized to a CM5 sensor chip (GE Healthcare). Serial dilutions of purified recombinant *Nanosota-1* drugs were injected at different concentrations: 320 nM – 10 nM for *Nanosota-1A*; 80 nM - 2.5 nM for *Nanosota-1B* and *Nanosota-1C*; 20 nM - 1.25 nM for *Nanosota-1C-Fc*. The resulting data were fit to a 1:1 binding model using Biacore Evaluation Software (GE Healthcare).

### Protein pull-down assay

Protein pull-down assay was performed using Immunoprecipitation kit (Invitrogen) as previously described (*20*). Briefly, 10 µl protein A beads were incubated with 1 µg SARS-CoV-2 RBD-Fc at room temperature for 1 hour. Then different amounts (7.04, 3.52. 1.76, 0.88, 0.44, 0.22, or 0 µg) of *Nanosota-1C* (with a C-terminal His tag) and 4 µg human ACE2 (with a C-terminal His tag) were added to the RBD-bound beads. After one-hour incubation at room temperature, the bound proteins were eluted using elution buffer (0.1 M glycine pH 2.7). The samples were then subjected to SDS-PAGE and analyzed through Western blot using an anti-His antibody.

### Gel filtration chromatography assay

Gel filtration chromatography assay was performed on a Superdex200 column. 500 µg human ACE2, 109 µg *Nanosota-1C* and 121 µg SARS-CoV-2 RBD were incubated together at room temperature for 30 min. The mixture was subjected to gel filtration chromatography. Samples from each peak off the column were then subjected to SDS-PAGE and analyzed through Coomassie blue staining.

### SARS-CoV-2 pseudovirus entry assay

The potency of *Nanosota-1* drugs in neutralizing SARS-CoV-2 pseudovirus entry was evaluated as previously described (*20, 22*). Briefly, HEK293T cells were co-transfected with a plasmid carrying an Env-defective, luciferase-expressing HIV-1 genome (pNL4-3.luc.R-E-) and pcDNA3.1(+) plasmid encoding SARS-CoV-2 spike protein. Pseudoviruses were collected 72 hours after transfection, incubated with individual drugs at different concentrations at 37°C for one hour, and then were used to enter HEK293T cells expressing human ACE2. After pseudoviruses and target cells were incubated together at 37°C for 6 hours, the medium was changed to fresh medium, followed by incubation of another 60 hours. Cells were then washed with PBS buffer and lysed. Aliquots of cell lysates were transferred to plates, followed by the addition of luciferase substrate. Relative light units (RLUs) were measured using an EnSpire plate reader (PerkinElmer). The efficacy of the drug was expressed as the concentration capable of neutralizing 50% of the entry efficiency (Neutralizing Dose 50 or ND_50_).

### SARS-CoV-2 plaque reduction neutralization test

The potency of *Nanosota-1* drugs in neutralizing authentic SARS-CoV-2 infections was evaluated using a SARS-CoV-2 plaque reduction neutralization test (PRNT) assay. Specifically, individual drugs were serially diluted in DMEM and mixed 1:1 with 80 pfu SARS-CoV-2 at 37°C for 1 hour. The mixtures were then added into Vero E6 cells at 37°C for an additional 45 minutes. After removing the culture medium, cells were overlaid with 0.6% agarose and cultured for 3 days. Plaques were visualized by 0.1% crystal violet staining. The efficacy of each drug was calculated and expressed as the concentration capable of reducing the number of virus plaques by 50% compared to control serum-exposed virus (i.e., ND_50_).

### SARS-CoV-2 challenge of hamsters

Equal sex Syrian hamsters (n=24) were obtained from Envigo (IN) and challenged via intranasal inoculation with SARS-CoV-2 (at a titer of 1 × 10^6^ Median Tissue Culture Infectious Dose or TCID_50_) in 100 µL DMEM (50 µL per nare). Sample size was comparable to previous animal challenge studies (*33*) and constrained by the availability of resources. At a sample size of 6 animals per group, G*Power analysis indicates that we can detect an effect size of 1.6 with a power of .80 (alpha = .05 one-tailed). Four groups of hamsters (n=6 each randomly assigned) were treated with *Nanosota-1C-Fc* via intraperitoneal injection at one of the following time points and dosages: (1) 24 hours pre-challenge at 20 mg/kg body weight of hamsters; (2) 4 hours post-challenge at 20 mg/kg body weight of hamsters; (3) 4 hours post-challenge at 10 mg/kg body weight of hamsters. Hamsters in the control (negative) group were administered PBS buffer 24 hours pre-challenge. An additional group was tested for a different hypothesis and the data were not included in the current study. Body weights were collected daily beginning prior to challenge. Nasal swabs were collected prior to challenge and additionally 1 day, 2 days, 3 days, 5 days and 10 days post-challenge for quantitative real-time RT-PCR (nasal swabs collected on day 2 and day 3 were lost due to Hurricane Laura). Hamsters were humanely euthanized 10 days post-challenge via overexposure to CO_2_. The lungs and bronchial tubes were collected and fixed in formalin for histopathological analysis. This experiment was performed in accordance with the guidelines set by the Institutional Animal Care and Use Committee at the University of Texas Medical Branch (UTMB).

### Half-life of Nanosota-1 drugs in mice

Male C57BL/6 mice (3 to 4 weeks old) (Envigo) were intravenously injected (tail-vein) with *Nanosota-1C* or *Nanosota-1C-Fc* (100 µg in 100 µl PBS buffer). At varying time points, mice were euthanized and whole blood was collected. Then sera were prepared through centrifugation of the whole blood at 1500xg for 10 min. The sera were then subjected to ELISA for evaluation of their SARS-CoV-2 RBD-binding capability.

### Biodistribution of Nanosota-1 drugs in mice

To evaluate the *in vivo* biodistribution of *Nanosota-1C-Fc* and *Nanosota-1C*, the nanobodies were labeled with Zirconium-89 [^89^Zr] and injected into male C57BL/6 mice (5 to 6 weeks old) (Envigo). Briefly, the nanobodies were first conjugated to the bifunctional chelator p-SCN-Bn-Deferoxamine (DFO, Macrocyclic) as previously described (*35*), and [^89^Zr] (University of Wisconsin Medical Physics Department) was then conjugated as previously described (*36*). [^89^Zr]-labeled nanobodies (1.05 MBq, 1-2 µg nanobody, 100 µl PBS) were intravenously injected (tail-vein). Mice were euthanized at different time points. Organs were collected and counted on an automatic gamma-counter (Hidex). The total number of counts per minute (cpm) for each organ or tissue was compared with a standard sample of known activity and mass. Count data were corrected to both background and decay. The percent injected dose per gram (%ID/g) was calculated by normalization to the total amount of activity injected into each mouse.

## Supplementary materials for

**Table S1.**
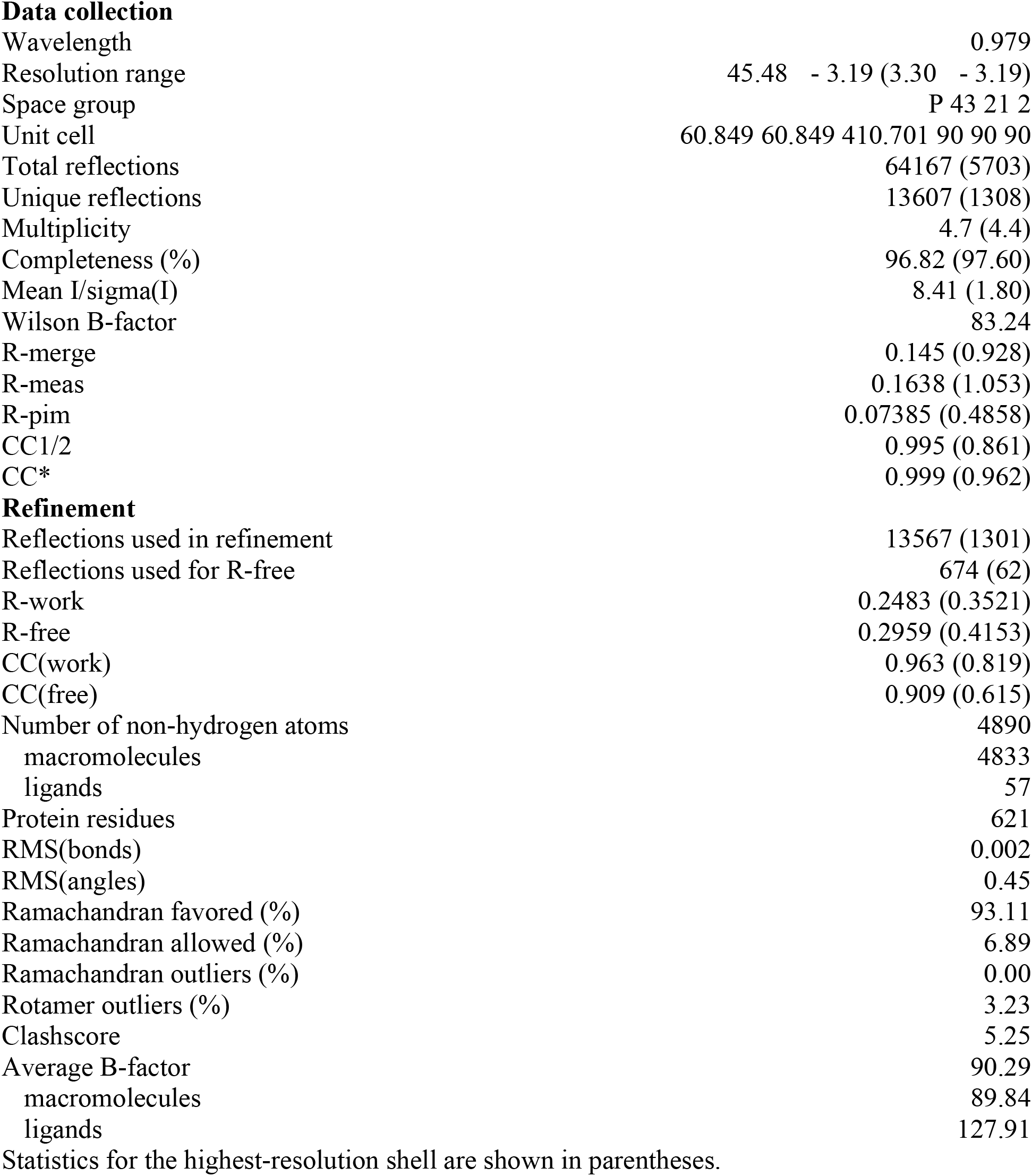
X-ray data collection and structure refinement statistics (SARS-CoV-2 RBD/*Nanosota-1C* complex)

|  |  |
| --- | --- |
| Wavelength | 0.979 |
| Resolution range | 45.48 - 3.19 (3.30 - 3.19) |
| Space group | P 43 21 2 |
| Unit cell | 60.849 60.849 410.701 90 90 90 |
| Total reflections | 64167 (5703) |
| Unique reflections | 13607 (1308) |
| Multiplicity | 4.7 (4.4) |
| Completeness (%) | 96.82 (97.60) |
| Mean I/sigma(I) | 8.41 (1.80) |
| Wilson B-factor | 83.24 |
| R-merge | 0.145 (0.928) |
| R-meas | 0.1638 (1.053) |
| R-pim | 0.07385 (0.4858) |
| CC1/2 | 0.995 (0.861) |
| CC* | 0.999 (0.962) |
| **Refinement** |  |
| Reflections used in refinement | 13567 (1301) |
| Reflections used for R-free | 674 (62) |
| R-work | 0.2483 (0.3521) |
| R-free | 0.2959 (0.4153) |
| CC(work) | 0.963 (0.819) |
| CC(free) | 0.909 (0.615) |
| Number of non-hydrogen atoms | 4890 |
| macromolecules | 4833 |
| ligands | 57 |
| Protein residues | 621 |
| RMS(bonds) | 0.002 |
| RMS(angles) | 0.45 |
| Ramachandran favored (%) | 93.11 |
| Ramachandran allowed (%) | 6.89 |
| Ramachandran outliers (%) | 0.00 |
| Rotamer outliers (%) | 3.23 |
| Clashscore | 5.25 |
| Average B-factor | 90.29 |
| macromolecules | 89.84 |
| ligands | 127.91 |

Statistics for the highest-resolution shell are shown in parentheses.

**Figure S1.**
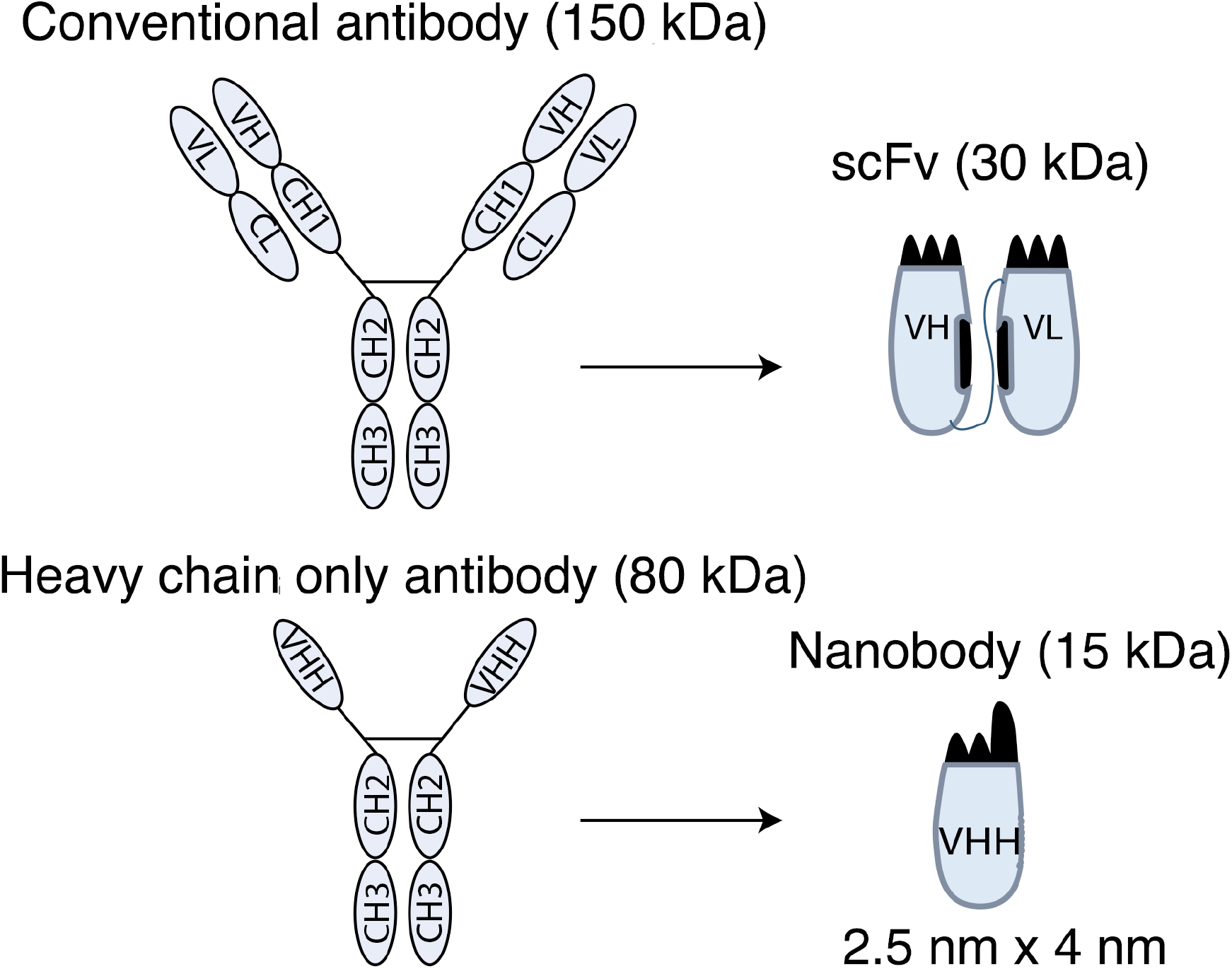
Schematic drawings of nanobodies and conventional antibodies. VH: variable domain of heavy chain. CH: constant domain of heavy chain. VL: variable domain of light chain. CL: constant domain of light chain. VHH: variable domain of heavy-chain only antibody. scFv: single-chain variable fragment.

**Figure S2.**
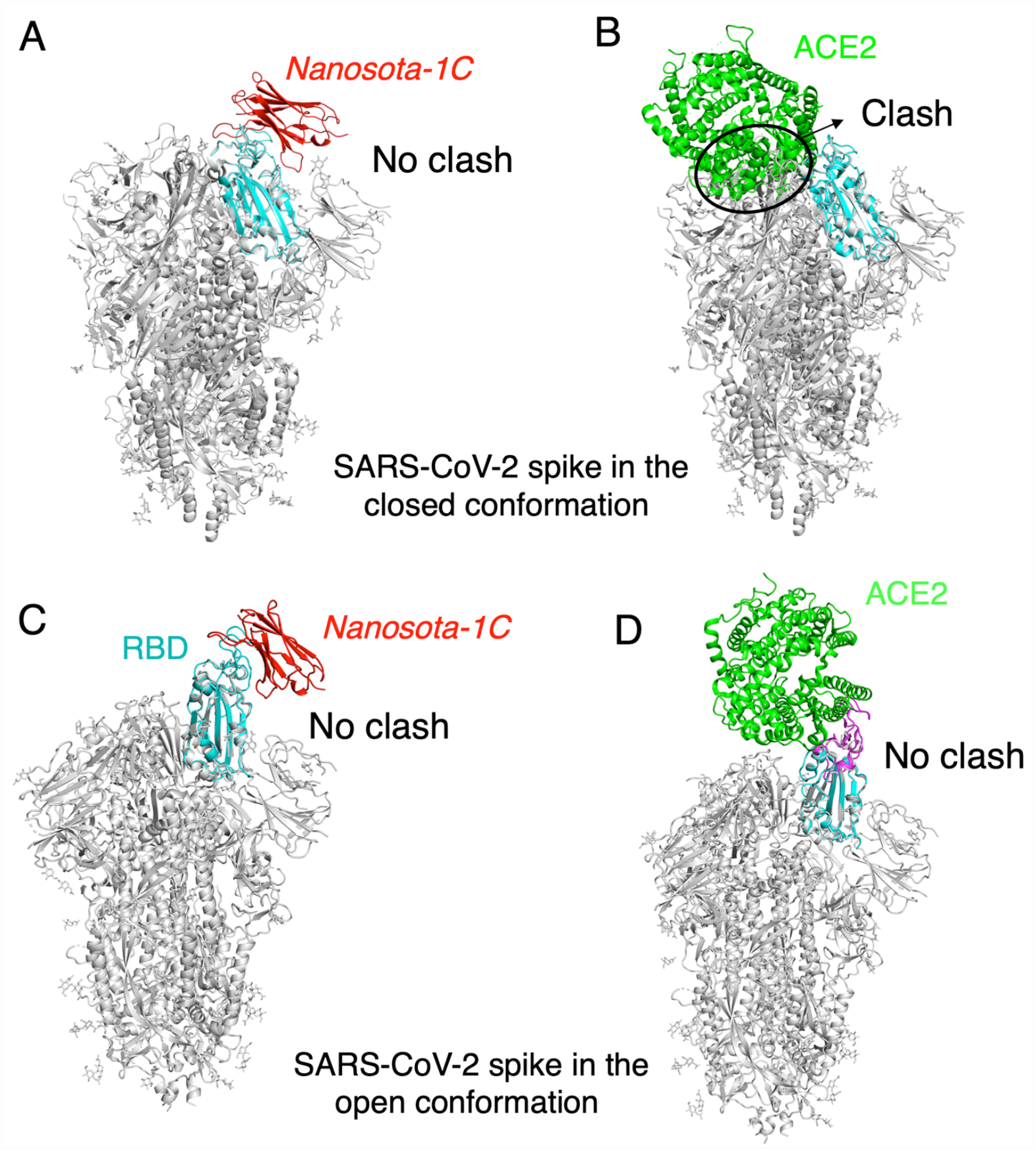
The binding of *Nanosota-1C* to SARS-CoV-2 spike protein in different conformations. (A) The binding of *Nanosota-1C* to the spike protein in the closed conformation. The structures of the RBD/*Nanosota-1C* complex and SARS-CoV-2 spike protein in the closed conformation (PDB: 6ZWV) were superimposed based on their common RBD structure (in cyan). *Nanosoto-1C* is in red. The rest of the spike protein is in gray. (B) The binding of ACE2 to the spike protein in the closed conformation. The structures of the RBD/ACE2 complex (PDB 6M0J) and SARS-CoV-2 spike protein in the closed conformation (PDB: 6ZWV) were superimposed based on their common RBD structure. ACE2 is in green. Clashes between ACE2 and the rest of the spike protein were circled. (C) The binding of *Nanosota-1C* to the spike protein in the open conformation (PDB: 6VSB). (D) The binding of ACE2 to the spike protein in the open conformation (PDB: 6VSB).

**Figure S3.**
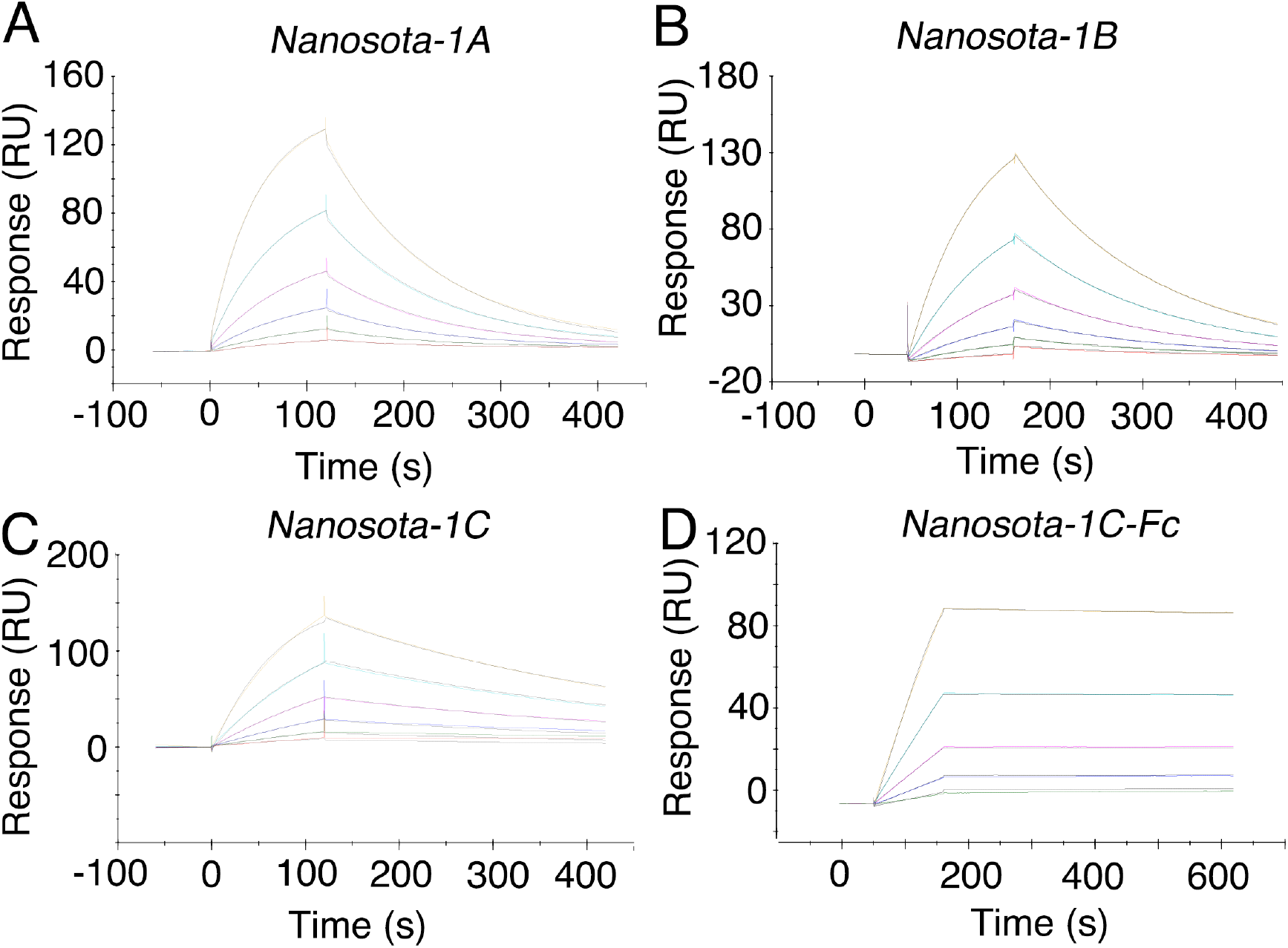
Measurement of the binding affinities between *Nanosota-1* drugs and SARS-CoV-2 RBD by surface plasmon resonance assay using Biacore. Purified recombinant SARS-CoV-2 RBD was covalently immobilized on a sensor chip through its amine groups. Purified recombinant nanobodies flowed over the RBD individually at one of five different concentrations. The resulting data were fit to a 1:1 binding model and the value of K_d_ was calculated for each nanobody. The assay was repeated three times (biological replication: new aliquots of proteins and new sensor chips were used for each repeat).

**Figure S4.**
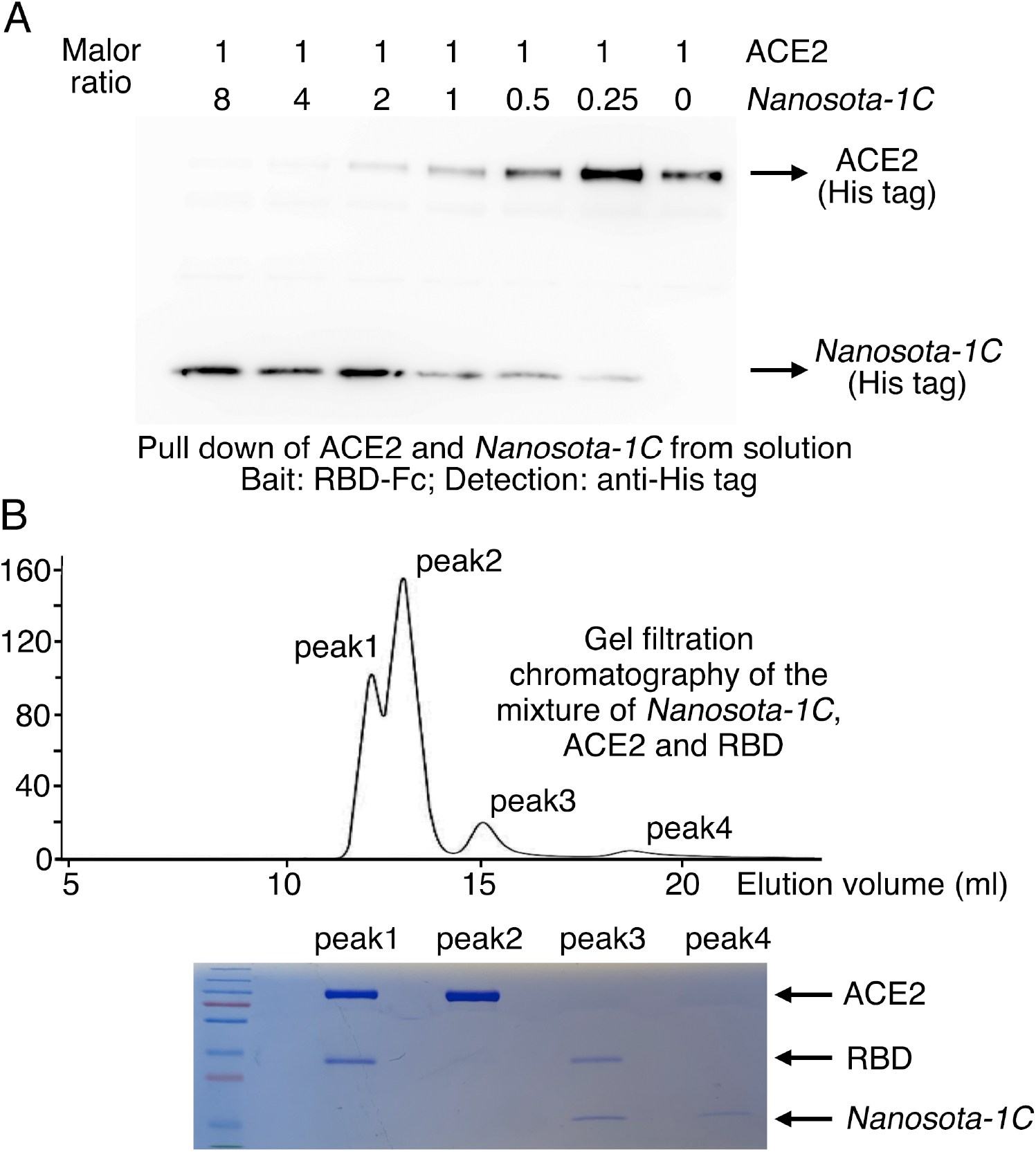
Binding interactions between *Nanosota-1* drugs and SARS-CoV-2 RBD. (A) Binding interactions between SARS-CoV-2 RBD, *Nanosota-1C*, and ACE2 as evaluated using a protein pull-down assay. Various concentrations of *Nanosota-1C* and a constant concentration of ACE2 (all His tagged) were combined in different molar ratios. SARS-CoV-2 RBD (Fc tagged) was used to pull down *Nanosota-1C* and ACE2. A western blot was used to detect the presence of *Nanosota-1C* and ACE2 following pull down by SARS-CoV-2 RBD. The assay was repeated three times (biological replication: new aliquots of proteins were used for each repeat). (B) Binding interactions between SARS-CoV-2 RBD, *Nanosota-1C*, and ACE2 as examined using gel filtration chromatography. *Nanosota-1C*, ACE2 and SARS-CoV-2 RBD (all His tagged) were mixed together in solution (both *Nanosota-1C* and ACE2 in molar excess of SARS-CoV-2 RBD) and purified using gel filtration chromatography. Protein components in each of the gel filtration chromatography peaks were analyzed with SDS-PAGE and stained by Coomassie blue. The assay was repeated three times (biological replication: new aliquots of proteins were used for each repeat).

**Figure S5.**
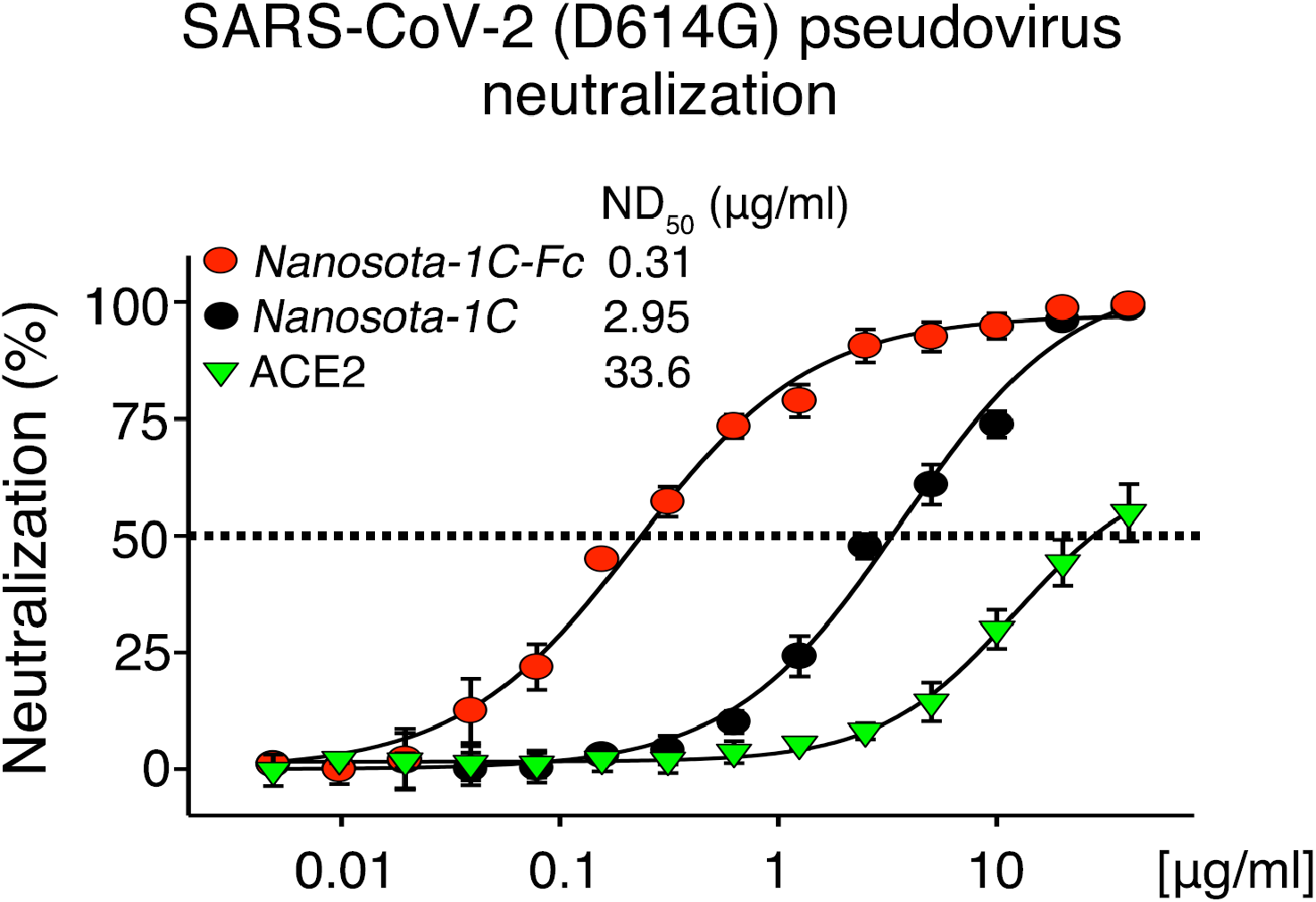
Neutralization of SARS-CoV-2 pseudovirus, which contains the D614G mutation in the spike protein, by *Nanosota-1* drugs. The procedure was the same as described in Fig. 3A, except that the mutant spike protein replaced the wild type spike protein. The assay was repeated three times (biological replication: new aliquots of pseudoviruses and cells were used for each repeat).

**Figure S6.**
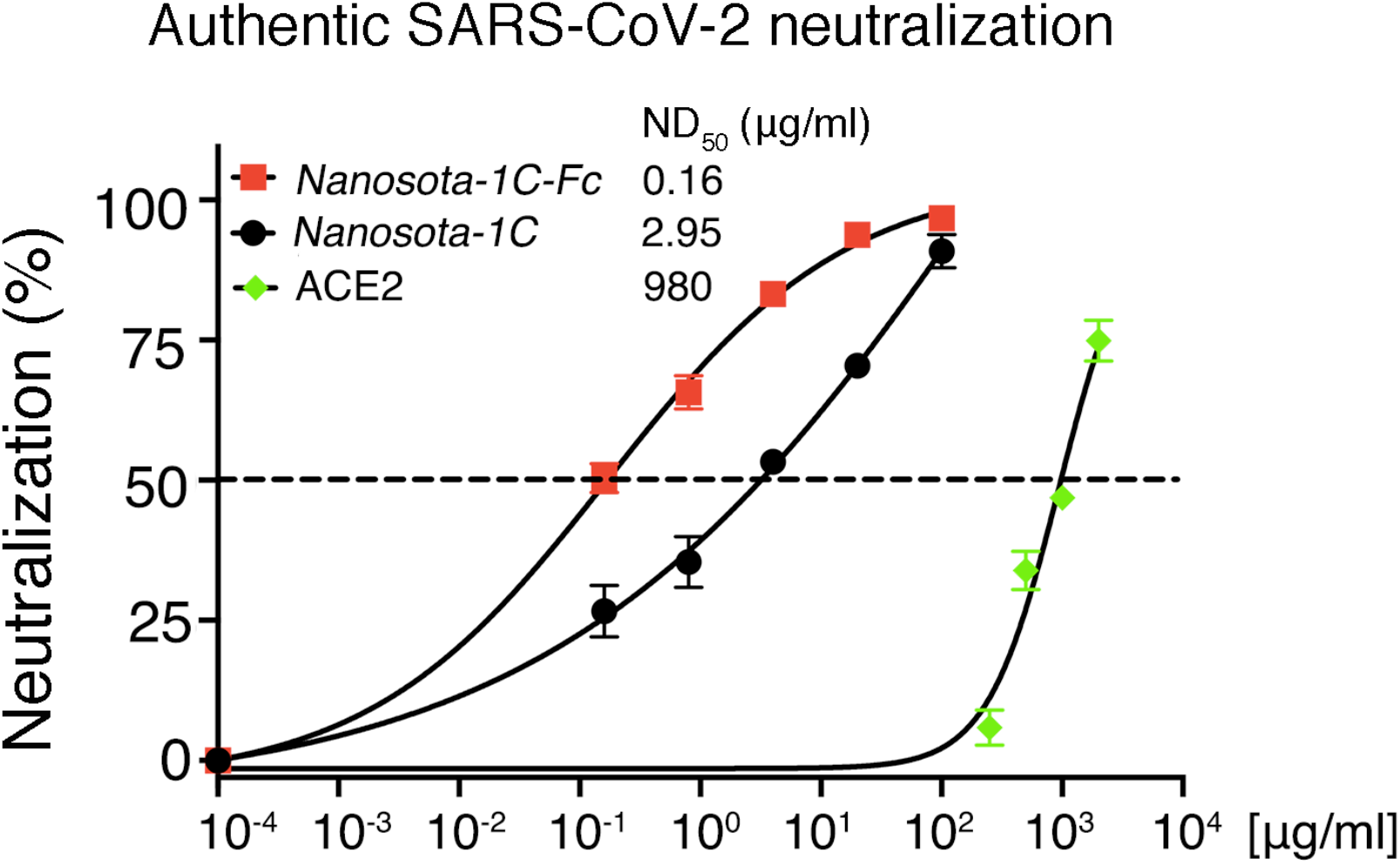
Detailed data on the neutralization of authentic SARS-CoV-2 infection of target cells by *Nanosota-1* drugs. Data are the mean ± SEM (n = 3). Nonlinear regression was performed using a log (inhibitor) versus normalized response curve and a variable slope model (R^2^ > 0.95 for all curves). The assay was repeated twice (biological replication: new aliquots of virus particles and cells were used for each repeat).

**Figure S7.**
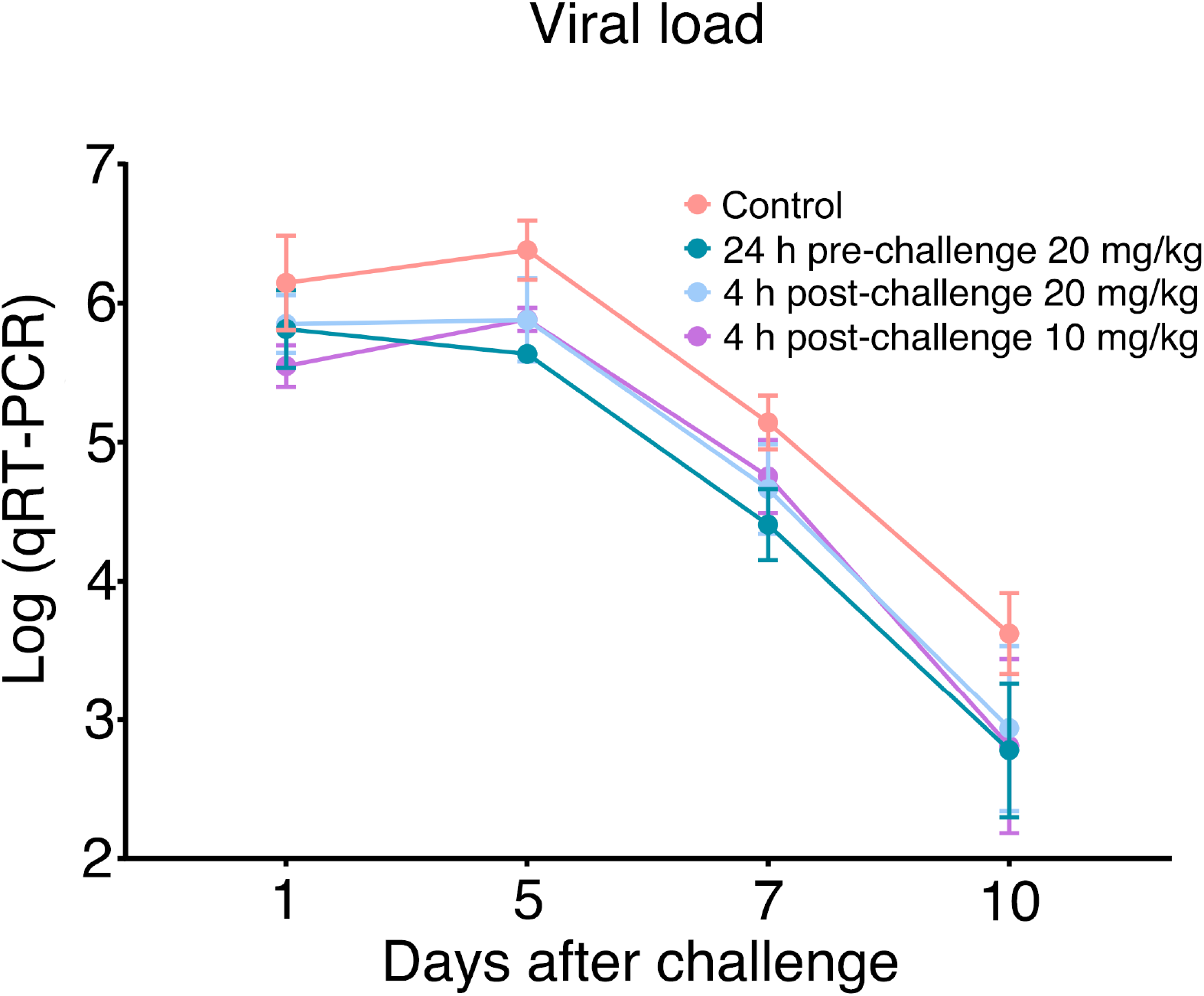
Additional data on the efficacy of *Nanosota-1* drugs in protecting hamsters from SARS-CoV-2 infections. Nasal swabs were collected from each hamster on days 1, 2, 3, 5, 7, and 10. Nasal swab samples from day 2 and day 3 were lost due to Hurricane Laura. qRT-PCR was performed to determine the virus loads in each of the samples. The qRT-PCR results are displayed on a log scale (since qRT-PCR amplifies signals on a log scale). Data are the mean ± SEM (n = 6). Missing data from one animal in the 4-hour post-challenge (10mg/kg) group on Day 7 were replaced by the average of that animal’s days 5 and 10 data. ANOVA analysis using group as a between-group factor and day (1, 5, 7, and 10) as a within-group factor revealed significant differences between the control group and each of the following groups: 24 hour pre-challenge (20 mg/kg) group (*F*(1, 10) = 6.02, *p* = .017, effect size *η*_*p*_^2^ = .38), 4 hour post-challenge (20 mg/kg) group (*F*(1, 10) = 5.38, *p* = .037, *η*_*p*_^2^ = .31), and 4 hour post-challenge (10 mg/kg) group (*F*(1, 10) = 3.40, *p* = .048, *η*_*p*_^2^ = .25). All *p-*values are one-tailed for directional tests.

**Figure S8.**
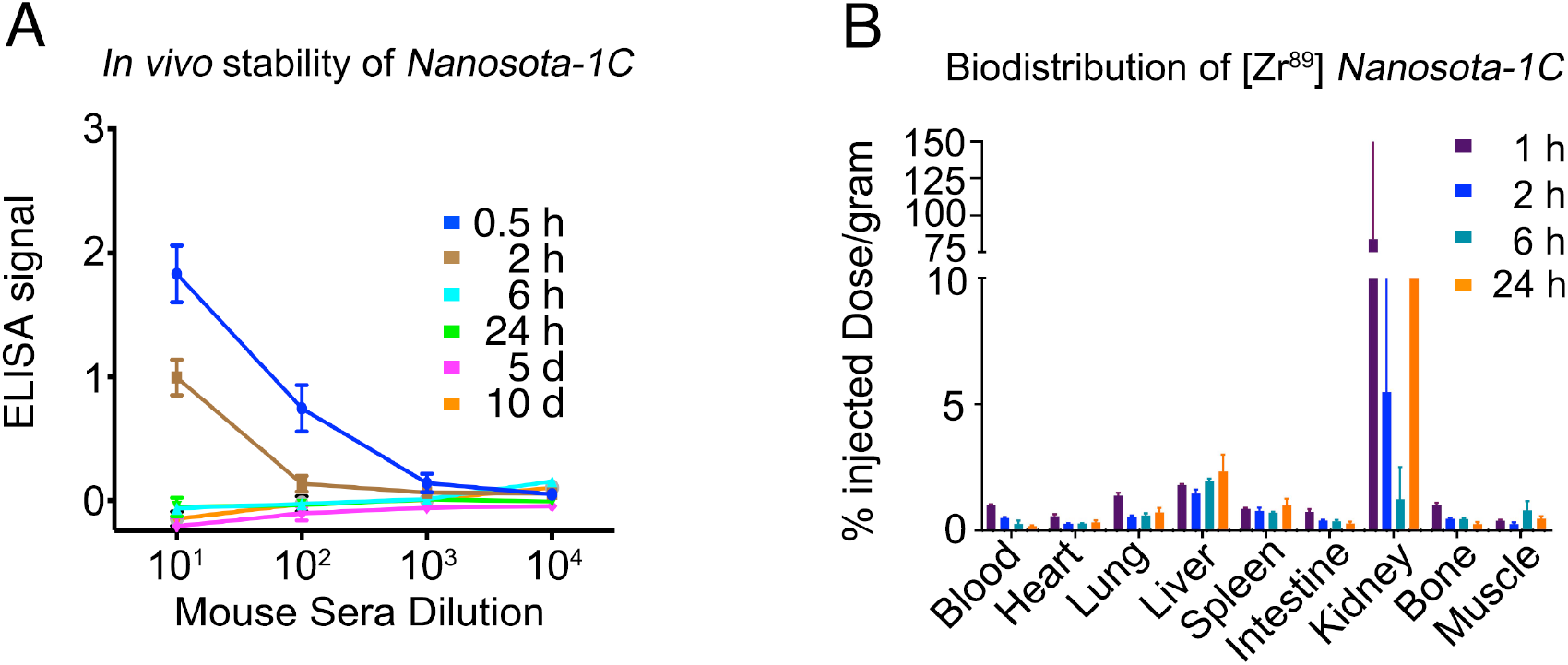
Pharmacokinetics of Nanosota-1C. *In vivo* stability and biodistribution of *Nanosota-1C* were measured in the same way as described in Fig. 5C and Fig. 5D, respectively, except that time points for *Nanosota-1C* differed from those for *Nanosota-1C-Fc* due to pharmacokinetic differences of the small molecular weight nanobody versus the larger Fc tagged nanobody.

